# Assessing dynamic functional connectivity in heterogeneous samples

**DOI:** 10.1101/118968

**Authors:** B. C. L. Lehmann, S. R. White, R. N. Henson, Cam-CAN, L. Geerligs

## Abstract

Several methods have been developed to measure dynamic functional connectivity (dFC) in fMRI data. These methods are often based on a sliding-window analysis, which aims to capture how the brain’s functional organization varies over the course of a scan. The aim of many studies is to compare dFC across groups, such as younger versus older people. However, spurious group differences in measured dFC may be caused by other sources of heterogeneity between people. For example, the shape of the haemodynamic response function (HRF) and levels of measurement noise have been found to vary with age. We use a generic simulation framework for fMRI data to investigate the effect of such heterogeneity on estimates of dFC. Our findings show that, despite no differences in true dFC, individual differences in measured dFC can result from other (non-dynamic) features of the data, such as differences in neural autocorrelation, HRF shape, connectivity strength and measurement noise. We also find that common dFC methods such as *k*-means and multilayer modularity approaches can detect spurious group differences in dynamic connectivity due to inappropriate setting of their hyperparameters. fMRI studies therefore need to consider alternative sources of heterogeneity across individuals before concluding differences in dFC.

## 1. Introduction

Brain connectivity can refer to a number of different types of relation between distinct regions in the brain. While structural connectivity refers to the anatomical links between brain regions, functional connectivity (FC) describes how activity in different regions is related over time. In functional magnetic resonance imaging (fMRI), this is commonly measured using the Pearson correlation between the fMRI time series in different brain regions (Biswal et al. (1995)). Typically, one would calculate the correlation between two time series over the course of the whole fMRI scan. However, this approach may represent an average across informative fluctuations in FC. Indeed, recent evidence suggests that even in task-free, resting-states these functional connections change over the course of a scan (Allen et al. (2012), Chang and Glover (2010), Kiviniemi et al. (2011)). Moreover, measures of this dynamic functional connectivity (dFC) have been used in an attempt to identify biomarkers for schizophrenia (Sakoğlu et al. (2010)) and Alzheimer’s disease (Jones et al. (2012)).

The most common way to measure dFC is to apply a sliding-window analysis (see Hutchison et al. (2013) for a review of dFC). Methods to analyse the changes in connectivity across windows vary in complexity. A simple approach characterises dFC as the standard deviation (SD) of the correlation values across time windows (Elton and Gao (2015)). Alternatively, one can pool data across individuals and use *k*-means clustering to identify recurring connectivity patterns (Allen et al. (2012)), or “FC states”. Another class of methods applies network theory on an individual level. The brain can be characterised as a complex graph, with distinct brain regions corresponding to nodes, and functional connections corresponding to edges between nodes (Bullmore and Sporns (2009)). Differences in the properties of the resulting graphs can then be used as measures of dFC (Bassett et al. (2011), Bassett et al. (2013b)). Crucially, both these classes of methods require a choice of hyperparameters, which have to be estimated from the data. For example, in a k-means analysis, the number of clusters *k* has to be prespecified.

Some of the challenges facing current methods, such as the choice of window width and the effect of data pre-processing, have already been discussed in Hutchison et al. (2013). Furthermore, (Shakil et al. 2016) have shown that the width and offset of windows can have an effect on the detection of FC state transitions and duration in a *k*-means analysis. In this paper we address an additional issue, namely unaccounted heterogeneity between individuals. While the aim of group studies is to detect heterogeneity in true dFC, other sources of heterogeneity may have an impact on estimated dFC. Heterogeneity can arise in a variety of ways. For example, Arbabshirani et al. (2014a) found the autocorrelation of fMRI time series within brain regions to differ between healthy brains and those with schizophrenia, and autocorrelation is known to affect estimation of cross-correlation (Arbabshirani et al. (2014b)). Although it is unclear whether this change in autocorrelation is due to neural or vascular factors, work with dynamic causal modeling has suggested that neural autocorrelation within some networks can vary between young and older participants (Tsvetanov et al. (2016)). Another example of heterogeneity is differences in the haemodynamic response function (HRF). The shape of the HRF, which can be modelled as a finite impulse response kernel, has been found to vary between healthy patients and patients with schizophrenia (Hanlon et al. (2016)) and also between age groups (Huettel et al. (2001), Aizenstein et al. (2004), D’Esposito et al. (1999)). Even non-neural physiological noise levels might differ across groups, owing for example to greater within-scan head movement in old relative to young subjects (Geerligs et al. (2015)).

**Figure 1:**
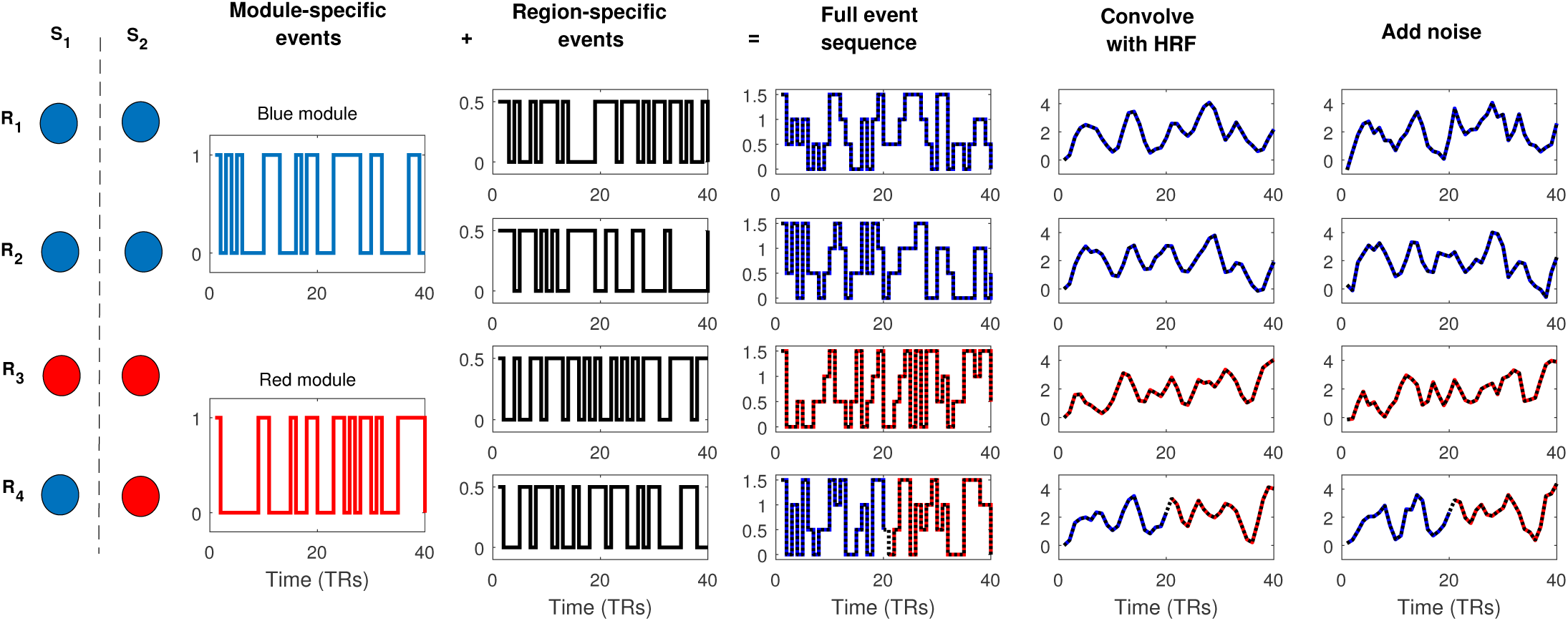
The generic simulation framework used to generate fMRI data for a brain consisting of four regions. State *i* is denoted by *S*_*i*_ and region *j* is denote by *R*_*j*_. Two brain regions are connected at any given time if they are in the same module, which are distinguished here by colours. These panels show data for 40 TRs: the first 20 TRs are spent in *S*_1_ while the last 20 TRs are spent in *S*_2_.

Here, we aim to investigate how unaccounted heterogeneity impacts estimates of dFC. Building on previous work by Allen et al. (2012), we designed a simulation framework to generate data from a dynamic connectivity structure based on FC states. We characterise FC states as time periods in which brain regions can be grouped into specific sets, or “modules”. In a given FC state, regions are considered connected if, and only if, they are in the same module. In this framework, changes in connectivity structure then correspond to FC state transitions. Data were generated to investigate the effect of individual differences in neural autocorrelation, HRF shape, connectivity strength and measurement noise on estimated dFC. We specifically used the case of aging to illustrate how plausible age-related sources of heterogeneity could impact dFC estimates. Furthermore, we varied the number of FC states and frequency of FC state transitions, in order to explore the effect of hyperparameter selection on the results of popular dFC methods such as the k-means method used by Allen et al. (2012) and the multilayer modularity approach of Bassett et al. (2011). Our analysis follows a typical sliding-window dFC pipeline based on a number of recent studies (e.g. Allen et al. (2012), Elton and Gao (2015), Sakoğlu et al. (2010)). Our findings show that group-level differences in neural autocorrelation, HRF shape, connectivity strength, measurement noise, number of FC states and frequencies of FC state transitions can lead to systematic differences in observed dFC between simulated fMRI time-series data.

## 2. Methods: simulation framework

To demonstrate some of the issues associated with assessing dFC in heterogeneous samples, we developed a simulation framework based on Allen et al. (2012). For each type of heterogeneity, we report the effects of changing only one parameter at a time, in order to isolate its relative impact on the analysis, though we consider interactions between parameters in the Supplementary Material. Thus, each source of heterogeneity corresponded to a change in a single step of the data generation process. To illustrate the effects of neural autocorrelation, HRF, connectivity strength and measurement noise, it was sufficient to simulate data from a model with only four regions of interest (ROIs) - Base Simulation 1. In order to analyse the impact of changes in both number of FC states and frequency of FC state transitions, we increased the number of ROIs to 32 to allow for a greater variety of states - Base Simulation 2. We now describe the simulation framework in detail, illustrated schematically in Figure 1, before outlining both Base Simulations and the specific variations for each source of heterogeneity.

We characterised FC states as time periods in which regions are partitioned into “modules”. For convenience, we denote state *i* as *S*_*i*_ and region *j* as *R*_*j*_. To simulate fMRI data for an individual, we first generated a FC state sequence to describe the changes in functional connectivity. This process is outlined in detail in the Base Simulations below. We used a sampling rate of TR = 2s and generated binary neural event sequences of length *T* = 360 TRs for each module and each region. The module-specific event sequences drive the connectivity structure: a module-specific event is one which occurs for all regions within that module in the current FC state. In contrast, region-specific events are those which occur for single regions only, independent of other regions, and thus correspond to neural noise.

More precisely, a module-specific event occurred in a module at an individual time point with probability *P*_*mod*_ = 0.5, independent of all other modules and time points. If a module has an event at time *t*, all regions within that module at time t have an event. For each region, we then superimposed a region-specific neural event sequence. A region-specific event occurred in a region at an individual time point with probability *P*_*reg*_ = 0.5, independent of all other regions and time points (except in Section 3.1, where we explored the effect of auto-correlated region-specific events). We fixed the amplitude of region-specific events to be *a*_*reg*_ = 1, and set the amplitude of the module-specific events to be *a*_*mod*_ = 2. The full event sequences were then convolved with a haemodynamic response function (HRF; kernel length = 16 TRs) using the SPM12 software (http://www.fil.ion.ucl.ac.uk/spm) to produce fMRI-like time series, which were rescaled to have a SD of 1. White noise with SD σ_*noise*_ = 0.2 was then added. Finally, a high-pass Butterworth filter removing frequencies below 0.033Hz was applied. This is based on the rule of thumb given by Leonardi and Ville (2015) which recommends removing frequency components below 1/*w*, where *w* is the window length in the sliding-window analysis.

### 2.1. Base Simulation 1 (4 ROIs)

In this setting, which corresponds to the framework illustrated in Figure 1, we restricted dynamics to two FC states, *S*_1_ and *S*_2_. *S*_1_ corresponded to the partition {1,1,2,1}, so that *R*_1_, *R*_2_, *R*_4_ were grouped into module 1, and *R*_3_ was grouped by itself into module 2, while *S*_2_ corresponded to the partition {1,1,2,2}. We fixed the FC state sequence such that each individual spent half of the time in *S*_1_ and then transitioned to *S*_2_. This allowed for the comparison of dFC between three types of region pairs: connected (within-module e.g. *R*_1_-*R*_2_), unconnected (between-module e.g. *R*_1_-*R*_3_), and a dynamic connection (within-module to between-module e.g. *R*_1_-*R*_4_). We then generated fMRI-like data using the simulation framework described above.

### 2.2. Base Simulation 2 (32 ROIs)

In this setting, we generated a total of 9 FC states, each consisting of a partition of the 32 ROIs into exactly 5 modules. For each FC state we generated a module label for each region from the numbers {1,…, 5} uniformly. If a FC state did not contain all 5 modules, we repeated this process. To ensure that no two FC states were too similar to each other, we computed the normalised mutual information (NMI) between each pair of state vectors, repeating the whole process if the maximum pairwise NMI exceeded 0.5. For each individual, we generated a random sequence of FC states under the assumption that a brain remained in a FC state for a fixed period of time before switching to any other FC state. Each FC state thus lasted a quarter of the total time period if three FC state transitions were specified, or half of the period if just one FC state transition was specified. We then generated fMRI-like data using the simulation framework described above.

## 3. Methods: specific simulations

For the first four simulations described here, we used Base Simulation 1 to generate the data. To measure dFC, we applied a sliding-window analysis. We used a tapered-cosine (Tukey) window of width *w* = 30 TRs with a total taper section of length 15 TRs. We slid the windows one time point at each step, yielding a total of 331 windows. We calculated pairwise Fisher-transformed Pearson correlation for each window and for each pair of regions. We then computed the SD of the time series of correlation values between each pair of regions. This measure is commonly defined as a proxy for dynamic functional connectivity. We also used the variance and the interquartile range of the correlation time series as alternative measures of dFC but these did not produce materially different results. For each set of parameters, we simulated 100 replicates in order to account for the randomness inherent in the data generation and also to assess the variability of our measure of dFC.

### 3.1. Neural autocorrelation

To investigate the effect of varying neural autocorrelation on the analysis of dFC, we used Base Simulation 1. To control the neural autocorrelation, we varied the generation of the region-specific neural event sequences. We modelled the binary sequences as Markov chains dependent on two parameters: the equilibrium probability of an event *π*_*reg*_ and the lag-1 autocorrelation *ρ*_*reg*_. The default value in Base Simulation 1 is the special case of *π*_*reg*_ = 0.5 and *ρ*_*reg*_ = 0, indicating no autocorrelation, and is the value used in later simulations. For the purposes of this simulation, we kept the equilibrium probability of an event fixed at *π*_*reg*_ = 0.5, thus ensuring that the expected number of events was constant at 180. We generated data for *ρ*_*reg*_ = −0.8, −0.7, …, 0.8 with the remainder of the simulation following the simulation framework described in section 2. We also performed this analysis with a range of values of *P*_*mod*_, *P*_*reg*_, *a*_*mod*_, and σ_*noise*_ and after prewhitening the data - see Figures S5 and S8 respectively.

Note that this region-specific signal can be considered a source of noise (as opposed to the module-specific events that drive the connectivity signals). The autocorrelation in this neural noise contributes to the temporal autocorrelation observed in the fMRI time series, which, once combined with the white noise measurement noise below, produces the AR(1)+white noise that characterises fMRI noise (at least after high-pass filtering; Friston et al. (2000)). Nonetheless, in real fMRI data, there are other sources of coloured noise, such as those induced by respiratory and cardiac signals, and by head-movement (see e.g. Woolrich et al. (2001)), which could also differ across groups.

**Figure 2:**
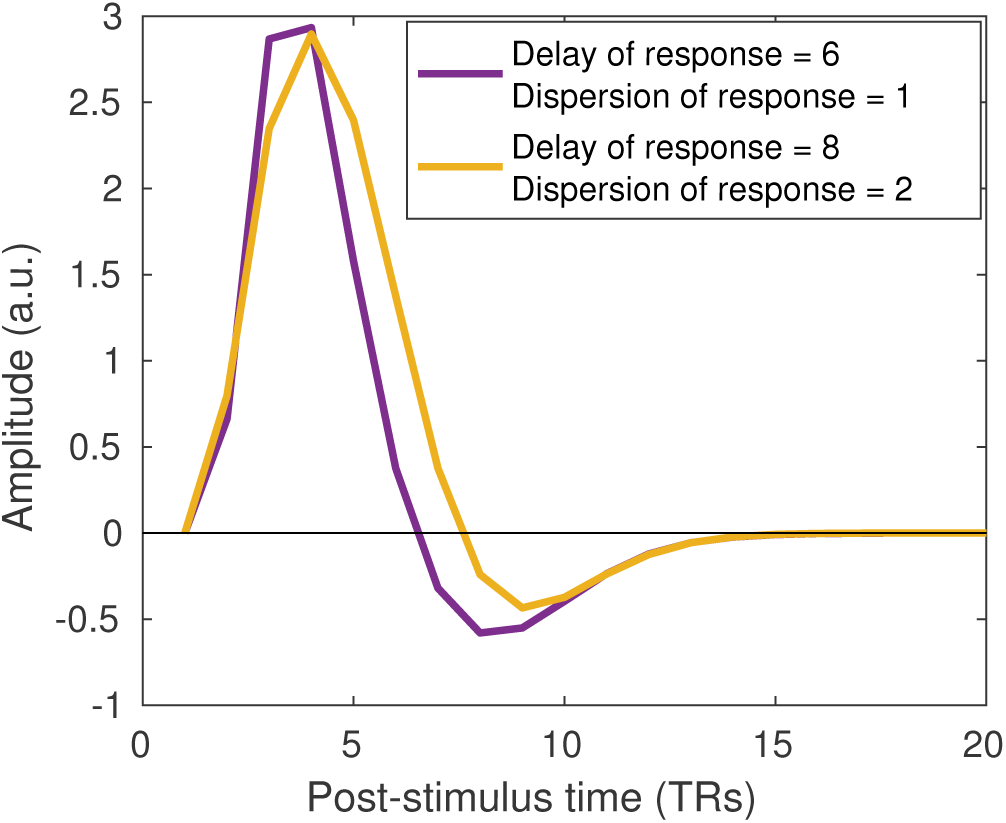
Two example haemodynamic response functions (HRF) based on different values for the dispersion of response and the delay of response. The purple line represents the default HRF used in the Base Simulations. We used a sampling rate of TR = 2s.

### 3.2. Haemodynamic response function

To demonstrate the impact of the HRF on dFC, we generated data with various HRFs, two of which are shown in Figure 2. We varied two of the HRF parameters, the dispersion of peak response (the width of the initial peak) and the delay of response, while the other 6 HRF parameters were held constant. In this simulation, we generated data using each HRF with peak dispersion σ_*HRF*_ = 0.6, 0.8,…, 2.4 and response delay τ_*HRF*_ = 5,5.2,…, 9s.

### 3.3. Connectivity strength

In our simulation framework, neural noise corresponds to the neural events that do not contribute to the connectivity structure. In other words, neural noise corresponds to the region-specific neural events, while the module-specific events are those which drive the connectivity structure. To investigate how connectivity strength affects dFC estimation, we varied the amplitude *a*_*mod*_ of the module-specific events relative to the region-specific events, which we fix to have amplitude *a*_*reg*_ = 1. In this simulation, we generated data for *a*_*mod*_ = 0.5, 1,…, 5. We also performed this analysis with a range of values of *P*_*mod*_, *P*_*reg*_, *ρ*_*reg*_, and σ_*noise*_ - see Figure S6.

### 3.4. Measurement noise

To investigate how measurement noise affects dFC estimation, we varied the amount of white noise added to the fMRI-like time series. We generated data with white noise of standard deviation σ_*noise*_ = 0, 0.1, …, 2.5 times that of the signal. We also performed this analysis with a range of values of *P*_*mod*_, *P*_*reg*_,*ρ*_*reg*_, and *a*_*mod*_ - see Figure S7.

### 3.5. k-means

A key assumption in many FC state-based methods is that there is a common set of FC states across individuals. In particular, it is assumed that, at rest, different participants cycle through a comparable set of brain states. In this simulation, we investigated what happens when this assumption does not hold. Specifically, we simulated 32-ROI fMRI data, using Base Simulation 2, from two groups of 50 people: group 1 could visit 9 distinct FC states, and group 2 could only visit 6 of these 9 FC states. Individuals in both groups experienced 3 FC state transitions in the course of a scan. FC state sequences for both groups were thus of length 4, with sequences for group 2 restricted to the FC states {1, …, 6}. For group 2, individuals were equally likely to be in each of the first six FC states. For group 1, in contrast, individuals were twice as likely to be in FC states 7,8 and 9 than in the first six FC states. The remainder of the simulation for each individual then followed the generic simulation framework.

We estimated correlation matrices for each position of the sliding-window analysis, as in the previous simulations, to produce 331 (Fisher-transformed) correlation matrices of size 32 × 32. The upper triangular part of this correlation matrix was vectorised to yield 331 correlation vectors of length 496 per subject. The *k*-means clustering was then performed on the set of all these vectors, pooled across subjects, with the ℓ_2_ norm as distance measure. Centroids were initialised using the *k*-means++ algorithm in Matlab and analysis was repeated 40 times with different initial centroids to avoid sub-optimal clusterings. We investigated the performance of the clustering with *k* = 1, …, 12 for the two groups separately and combined.

For each *k*, the algorithm returned a sequence of FC state labels for each subject, and the centroid of each of the *k* FC states. As the recovered FC state labels are arbitrary, the labels do not necessarily match those of the true FC states. While one can permute the recovered FC state labels to maximise the overlap with the true FC state sequence, this becomes more difficult with multiple subjects. Additionally, a simple relabelling does not take into account that incorrectly labelled FC states are not equally wrong. For example, if in two distinct windows, a subject is in FC state 1, but a k-means analysis recovers FC states 2 and 3 respectively (after relabelling), it may be the case that FC state 2 is closer to FC state 1 than FC state 3.

To circumvent the mislabelling and to enable comparisons of performance across different values of *k*, we replaced FC state labels by correlation matrices. For the recovered FC state sequences, we used the corresponding FC state centroid as calculated by the *k*-means algorithm. For the true sequences, we replaced a FC state label by a ‘true’ correlation matrix for that state. Recall that, in the 360 TRs simulated, a brain experienced 3 FC state transitions so that each FC state lasted 90 TRs. For each FC state, we first calculated the correlation matrix for each window of width 90 TRs in which all time points are in that FC state. We then took the ‘true’ correlation matrix as the average of all the corresponding correlation matrices in the same FC state across all subjects.

At each time point, we then computed the centroid error as the ℓ_2_ distance from the ‘true’ correlation matrix to the centroid of the recovered FC state at that time. We thus used two measures of performance: the average number of detected FC state changes across subjects, and the mean centroid error across all time points and subjects.

Note that, in typical task-free fMRI analyses, we do not know the ground truth. In our simulation, however, we assumed that the states of all participants were drawn from a larger common pool of states. The *k*-means algorithm identifies comparable connectivity patterns and groups them into states. It does not take into account the order in which the states occur for an individual and so a state can occur at different times for different participants. This allowed us to cluster across individuals and time points.

### 3.6. Multilayer Modularity

In many studies of dFC, a question of interest is whether groups differ in the degree to which connections between regions are static versus dynamic. Here, we examined how the frequency of state transitions could be detected using a multilayer moduarity algorithm, and how the choice of parameters affected these results. To this end, we simulated 32-ROI data, using Base Simulation 2, from two groups of 50 people: individuals in group 1 experienced 3 FC state transitions, and group 2 experienced just 1 FC state transition in the course of a scan. FC state sequences for group 1 were thus of length 4, while FC state sequences for group 2 were of length 2. All individuals could visit the same 9 FC states. The remainder of the simulation for each individual then followed the simulation framework described in Section 2. We applied the same sliding-window analysis as in the previous simulations, again using a window of width *w* = 30, calculating pairwise (Fisher-transformed) Pearson correlation for each window and for each pair of regions. In this case, however, we slid windows in steps of 30 TRs (instead of 1 TR) resulting 12 non-overlapping windows of width 30 TRs. This is based on the multilayer-modularity approach used by Bassett et al. (2011).

In the multilayer-modularity approach, the brain is characterised as a multilayer network with nodes corresponding to brain regions in different windows, or “layers”. The nonoverlapping sliding-window analysis yields a correlation matrix *A* with *A*_*ijl*_ corresponding to the correlation between regions *R*_*i*_ and *R*_*j*_ in window *l*. For each partitioning of regions into modules, the following multilayer modularity index is defined as a measure of the quality of the partition:

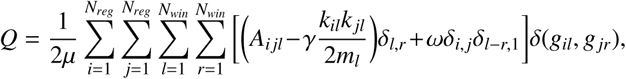
where *γ* and *ω* are hyperparameters, *N*_*reg*_ is the number of regions, *N*_*win*_ is the number of windows, *g*_*il*_ is the module assignment of region *R*_*i*_ in window *l, k*_*il*_ = Σ *jA_i_j_l_*, 2*m_l_* = Σ*_i_jA_i_j_l_*, and 2*μ* = Σ*j_l_*(*kj_l_* + Σ*_r_* ωδ_*l*−*r*,1_). The *δ* function is defined such that *δ*_*i, j*_ ≡ *δ*(*i, j*) = 1 if *i* = *j* and is equal to 0 otherwise. In our simulation, *N_reg_* = 32 and *N_win_* = 12. The regions can then be partitioned into modules by attempting to maximise the modularity index *Q* using a generalised Louvain algorithm (Mucha et al. (2010)).

We investigated the effect of varying the hyperparameters *γ* and ω on the accuracy of the subsequent partitions. Broadly speaking, *γ* controls the resolution of the partitioning within layers so that a high value of *γ* encourages regions to be grouped into smaller modules, thus increasing the total number of modules. On the other hand, ω influences the ‘stickiness’ of module assignments between consecutive layers. Thus, a high value of *ω* encourages fewer module changes for an individual region.

To assess the performance of the algorithm, we used two error measures. Firstly, we created “incidence matrices” of size 32 × 32 × 12, which contained an entry of 1 if the corresponding pair of regions had the same module label during a given window, and 0 otherwise. We then calculated the error in connectivity structure as the Hamming distance of the recovered incidence matrix (based on the multilayer modularity partitioning) to the true incidence matrix (based on the original module structure) at that time. The Hamming distance between two matrices of the same size is given by the number of elements at which the matrices differ. Secondly, we computed the mean flexibility for each subject, as defined by Bassett et al. (2011). The flexibility of a region is calculated by the number of times the region changes module assignment divided by the total possible number of module changes. The mean flexibility is then given by the mean region flexibility over all 32 regions. Note that the expected mean flexibility for a young individual in our simulation is approximately 4/55 (1 state change out of a possible 11 with the probability of a region changing module at a state change of approximately 0.8), compared to 12/55 for an old individual. For each subject, we ran the algorithm for *γ* = 0.75, 1,…, 2.5 and *ω* = 0.25, 0.5,…, 4.5.

## 4. Results: issues and limitations

We first investigate whether unaccounted heterogeneity can impact estimates of dFC. Here we examine four sources of heterogeneity: 1) individual differences in neural autocorrelation, 2) shape of the HRF, 3) connectivity strength and 4) measurement noise. We consider a simple, yet common, measure of dynamic connectivity, namely the standard deviation (SD) of correlation values across sliding windows. We calculated this measure for three types of true connectivity: 1) static, positive connections between regions within the same module, 2) static, zero connections between regions in different modules, and 3) dynamic changes between positive and zero connections when a region switched from being in the same module to being in a different module as another region (see Section 2). As expected, across all the simulations, estimated dFC for the dynamic connections is higher than the estimated dFC for both types of static connections.

### 4.1. Neural autocorrelation

In this simulation, we investigated the association between neural autocorrelation and estimated dFC, making sure that differences in neural autocorrelation were not associated with differences in ‘true’ dynamic connectivity. This was achieved by varying the autocorrelation *ρ*_*reg*_ of *region-specific* events but keeping the autocorrelation of the module-specific events fixed at zero. Recall that region-specific events are generated independently of the connectivity structure so, under our simulation framework, changing them should have no effect on dFC. Note that the underlying dFC structure is held constant across all iterations. We estimated dFC for three types of connection: a static, positive connection, a static zero connection, and a dynamic connection (from positive to zero) for *ρ*_*reg*_ = −0.8, −0.6,…, 0.8.

**Figure 3:**
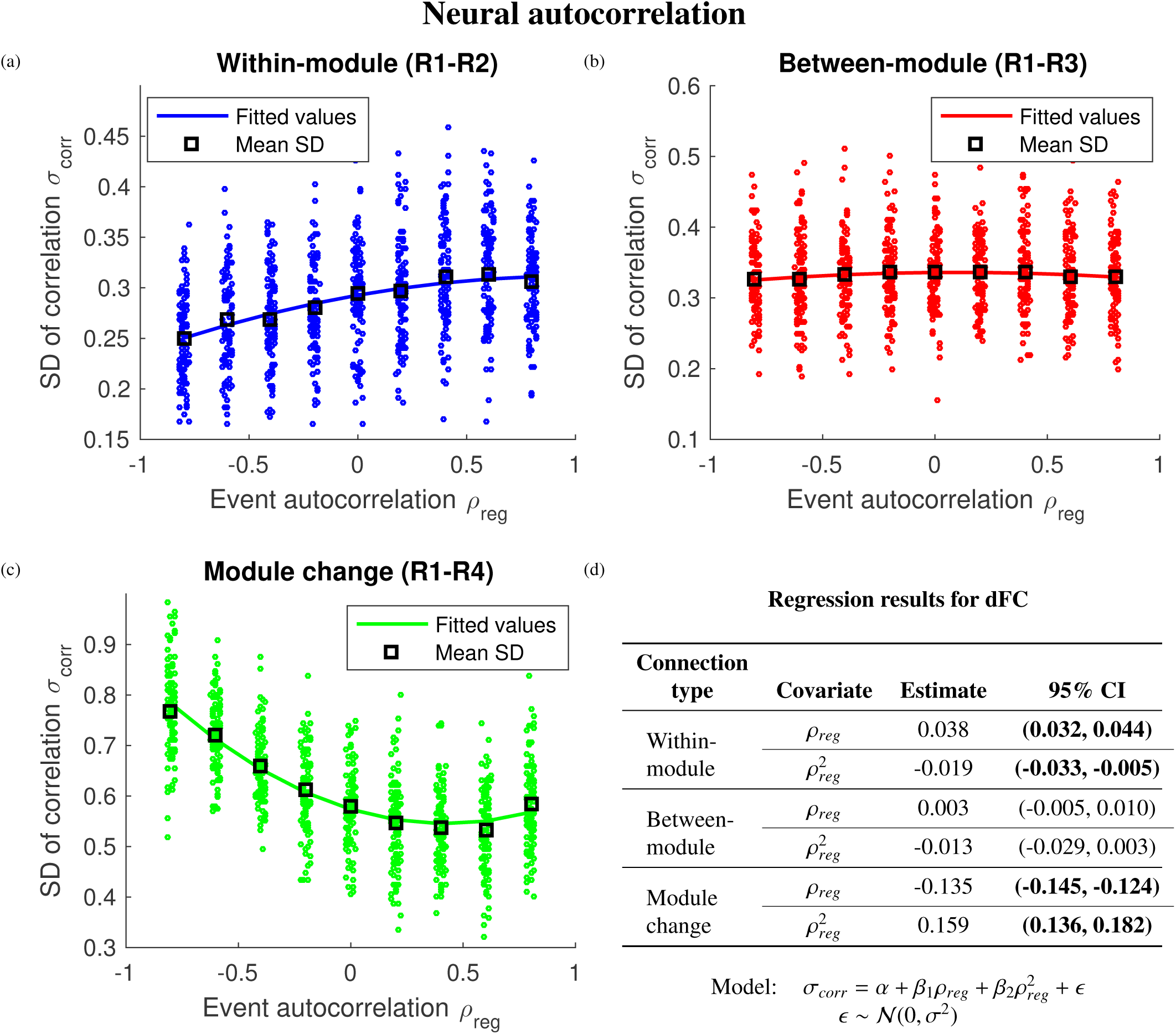
The impact of neural autocorrelation on estimated dFC, measured by the SD of the correlation time series, between (a) statically connected regions (within-module), (b) unconnected regions (between-module), and (c) dynamically connected regions (module change), with (d) the results of a multiple regression. R_*i*_ refers to Region i. Region pair R1-R4 has a dynamic connection so the true dFC should be higher than region pairs R1-R2 and R1-R3, which have a static connection. Estimated dFC between statically connected (within-module) regions increased with neural autocorrelation, while estimated dFC for dynamically connected regions (module change) decreased with increased neural autocorrelation. The neural autocorrelation is varied independently of the underlying connectivity structure, so changing it should have no effect on dFC. The multiple regression assesses the impact of neural autocorrelation on estimated dFC with a statistically significant effect indicated by a 95% confidence interval (CI) in bold type. The solid lines in (a-c) correspond to the fitted values of the multiple regression.

Figure 3 illustrates the impact of neural autocorrelation on estimated dFC. Although Figure 3b shows that estimated dFC for the unconnected regions remains unaffected by changes in neural autocorrelation, Figure 3a demonstrates that estimated dFC between the positively connected regions is higher as neural autocorrelation increases. In contrast, Figure 3c shows that estimated dFC decreased between the dynamically-connected regions as the autocorrelation increased. In other words, the ability to detect a difference in dFC between dynamically connected and statically connected regions decreases with higher levels of neural autocorrelation. These observations are supported by the multiple regression in Figure 3d: neural autocorrelation had a statistically significant effect on estimated dFC for the positively connected and dynamically connected regions, but not for the unconnected regions. These results suggest that observed dFC may vary substantially due to differences in neural autocorrelation, even though the true dFC was identical across individuals.

### 4.2. Haemodynamic response function

It has been shown that the haemodynamic response function (HRF) varies between different ages (Huettel et al. (2001), Aizenstein et al. (2004), D’Esposito et al. (1999)) and disease states (Hanlon et al. (2016)). To illustrate the effect of the HRF on observed dFC, we altered the HRF by varying the dispersion of the response σ_*HRF*_ and the delay of response τ_*hrf*_. Note that the underlying dFC structure is held constant across all iterations. To measure dFC, we calculated the mean SD of the sliding-window correlation time series for the three types of region pairs (positively connected, unconnected, and dynamically connected) for σ_*HRF*_ = 0.6, 0.8, …, 2.4 and τ_*hrf*_ = 5, 5.2, …, 9s.

Figures 4a and 4b show that a more temporally dispersed HRF (as often observed for older individuals) resulted in increased dFC between regions, when in truth the connectivity remained constant. Increases in both the dispersion of peak response σ_*HRF*_ and the delay of response τ_*HRF*_ resulted in increased estimated dFC. Figure 4c shows a similar effect for regions with a dynamic connection though, in this case, the increase was not as marked. Figure 4d compares the observed dFC for two of these HRFs. Individuals in Group 1 had a HRF with peak dispersion 1 and a response delay of 6s (purple line in Figure 2), which might represent a younger sample. In contrast, individuals in Group 2 had a HRF with peak dispersion 2 and a response delay of 8s (yellow line in Figure 2), which might represent an older sample. For the static connections we see that, even though the true dFC was the same across groups, the observed dFC varied substantially between groups due to the shape of the HRF. These observations are supported by the multiple regression results in Figure 4e: both the dispersion of peak response σ_*HRF*_ and the delay of response τ_*hrf*_ had statistically significant effects on estimated dFC for the statically connected regions, but only the delay of response τ_*hrf*_ had a statistically significant impact for the dynamically connected regions.

### 4.3. Connectivity strength

In our simulation framework, connectivity strength corresponds to the amplitude of the module-specific events, since these drive the connectivity structure. In this simulation, we investigated the association between connectivity strength and estimated dFC. This was done by varying the amplitude of module-specific events *a_mod_* while keeping the amplitude of the region-specific events fixed (i.e. the size of the connectivity “signal” versus region-specific neural “noise”). Note that the underlying dFC structure is held constant across all iterations. To measure dFC, we calculated the mean SD of the sliding-window correlation time series for the three types of region pairs (positively connected, unconnected, and dynamically connected) for *a*_*mod*_ = 0.5, 1, …, 5.

Figure 5 illustrates the impact of connectivity strength on estimated dFC. As expected, Figure 5b shows that estimated dFC for the unconnected regions remains unaffected by changes in connectivity strength. However, Figure 5a demonstrates that estimated dFC between the positively connected regions is moderately lower as connectivity strength increases. In contrast, Figure 5c shows that estimated dFC increased between the dynamically-connected regions as connectivity strength increased. In particular, the ability to detect a difference in dFC between dynamically connected and statically connected regions decreases with lower connectivity strength. These observations are supported by the multiple regression results in Figure 5d: connectivity strength had a statistically significant effect on estimated dFC for the positively connected and dynamically connected regions, but not for the unconnected regions. These results suggest that observed dFC may vary substantially due to differences in connectivity strength, even though the true dFC was identical across individuals.

### 4.4. Measurement noise

We also investigated the effects of varying amounts of measurement noise on observed connectivity dynamics by generating data with white noise of SD σ_*noise*_ = 0,0.1,…, 2.5. Recall that the signal was rescaled to have SD 1, resulting in noise-to-signal ratios equal to σ_*noise*_ (ignoring neural noise).

Figure 6 shows the effect of varying measurement noise on the standard deviation of correlation. Figures 6a, 6b, and 6c show that for all three types of region pair, increasing the amount of measurement noise resulted in decreased observed dFC. This is supported by the multiple regression results in Figure 6d: measurement noise had a statistically significant effect on estimated dFC for all three connection types. Thus, noisier data resulted in lower estimated dFC even for the pair of ROIs that experienced a true change in connectivity. When the amplitude reached a certain threshold (σ_*noise*_ > 2.0), the white noise dominated the underlying fMRI signal, resulting in a mean dFC of 0.2 for all three types of connectivity.

In the Supplementary Material, we demonstrate that the effects of neural autocorrelation, connectivity strength and measurement noise on estimated dFC persist across a range of parameter values (see Figures S5, S6 and S7 respectively).

### 4.5. k-means

In the previous sections, we demonstrated that group differences in connectivity dynamics, as measured with simple sliding window approaches, can be due to other factors such neural autocorrelation. However, even when such differences do not exist, some common dFC methods may still detect artifactual group differences in dFC owing to unaccounted heterogeneity in the dynamic connectivity structure.

To investigate the effect of heterogeneity in the number of FC states attainable, we generated a set of data for two groups of 50 individuals: those in G1 could reach 9 FC states, and those in G2 could reach only 6 of these 9 FC states (see Section 3.5). If these FC states can be recovered accurately from the data, then a simple measure of dFC is the number of FC state transitions that occur. Importantly, in the simulations, the number of such transitions was identical across groups, namely three.

**Figure 4:**
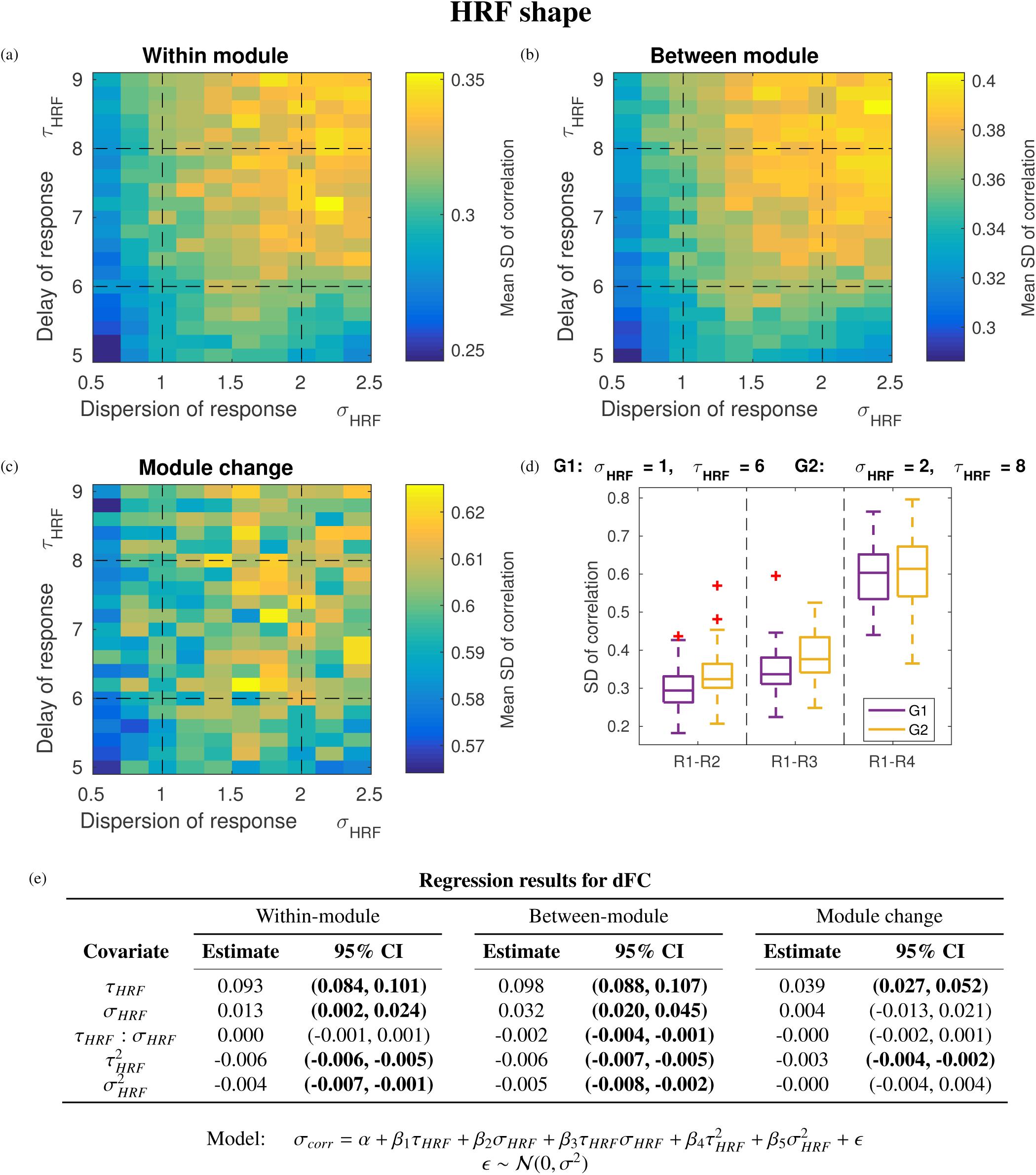
The impact of HRF shape on estimated dFC, measured by the SD of the correlation time series, between (a) statically connected regions (within-module), (b) unconnected regions (between-module), and (c) dynamically connected regions (module change), with (d) a comparison of estimated dFC for two groups and (e) a summary of results. *Ri* refers to Region *i*. The HRF was altered by varying two parameters: the dispersion of response σ_*HRF*_ and the delay of response τ_*HRF*_. Individuals in G1 had an HRF with σ_*HRF*_ = 1 and τ_HRF_ = 6 while individuals in G2 had an HRF with σ_*HRF*_ = 2 and τ_*HRF*_ = 8. Each square in (a-c) corresponds to the mean (across individuals) estimated dFC for a pair of parameter values of (σ_*HRF*_, τ_*HRF*_), with yellow indicating higher dFC. The dashed lines in (a-c) indicate the parameter values of the two groups compared in (d), whose HRFs are shown in Figure 2. A more dispersed HRF increased estimated dFC between all three types of region pairs, despite all individuals having identical dFC structure. We observed higher variability in dFC for the statically connected regions (within-module) as the dispersion and delay of the response increased. We saw a similar effect for the dynamically connected regions (module change), though this was less pronounced. While the box plots in (d) illustrate a single comparison of two groups, the multiple regression results in (e) summarise the impact of the two HRF parameters on estimated dFC with a statistically significant effect indicated by a 95% confidence interval (CI) in bold type.

**Figure 5:**
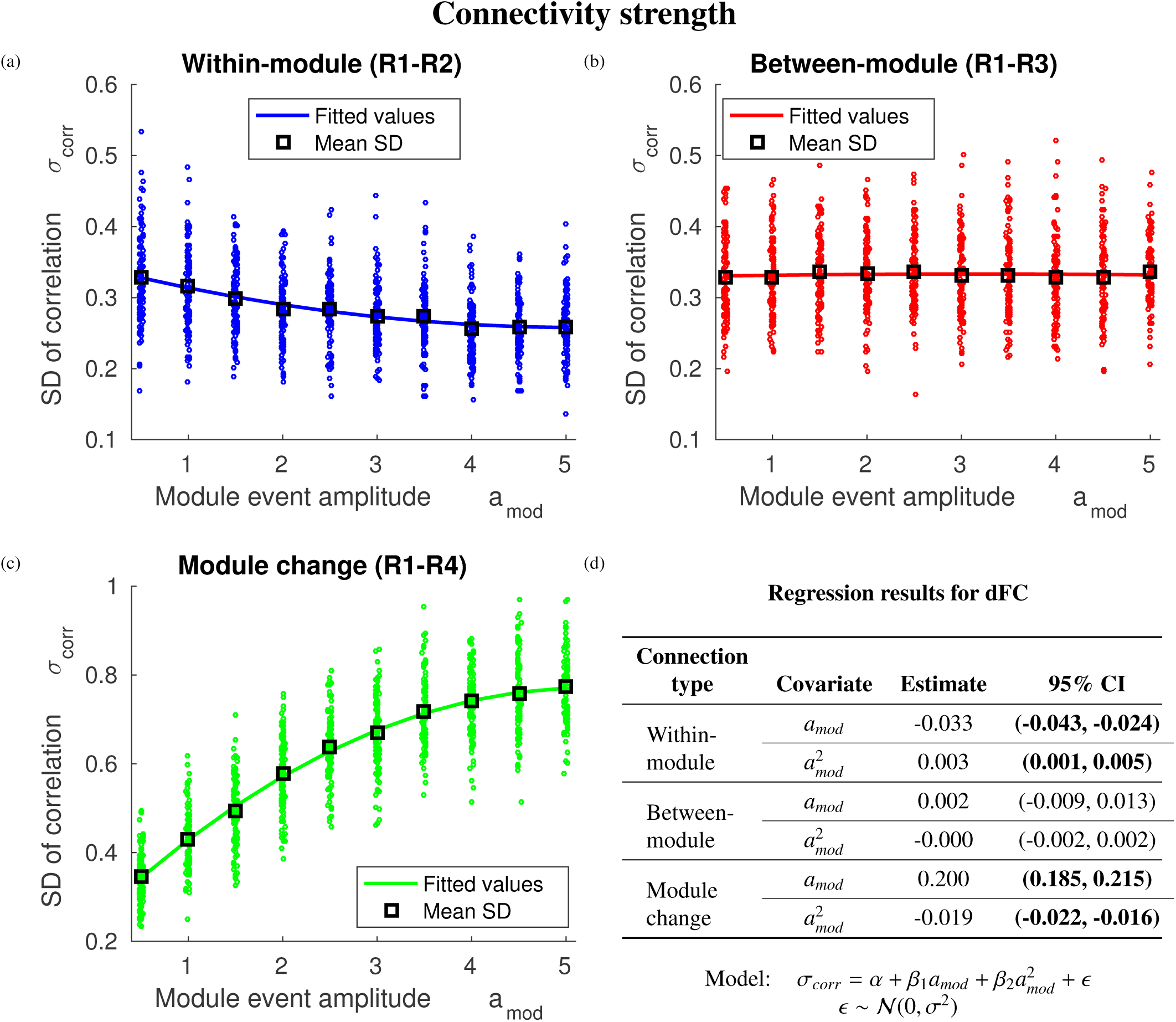
The impact of connectivity strength on estimated dFC, measured by the SD of the correlation time series, between (a) statically connected regions (within-module), (b) unconnected regions (between-module), and (c) dynamically connected regions (module change), with (d) the results of a multiple regression. R_*i*_ refers to Region *i*. Region pair R1-R4 has a dynamic connection so the true dFC should be higher than region pairs R1-R2 and R1-R3, which have a static connection. Increased amplitude of module-specific events resulted in decreased observed dFC for the positive static connected pair (within-module) but increased dFC for the dynamically connected pair (module change). The effect on the unconnected pair (between-module) was small. The connectivity strength, corresponding to the amplitude of module-specific events, is varied independently of the underlying connectivity structure, so changing it should have no effect on dFC. The multiple regression assesses the impact of connectivity strength on estimated dFC with a statistically significant effect indicated by a 95% confidence interval (CI) in bold type. The solid lines in (a-c) correspond to the fitted values of the multiple regression.

We used a *k*-means cluster analysis on the correlation matrices in an attempt to recover the underlying FC states. Figure 7 illustrates the performance of the *k*-means analysis for values of the hyperparameter *k* = 1, …, 12. We ran the analysis for the two groups separately, and also for all 100 individuals together. Here, we perform the analysis with sliding-window width *w* = 30, though window widths *w* = 60, 90, 120 TRs, yielded broadly similar results (see Figures S1, S2 and S3). Further, Figure S4 shows that, when *k* is correctly estimated or only slightly misspecified, it becomes more difficult to estimate the states correctly as window length increases.

Figure 7a shows that when *k* < 9, the typical (combined) *k*-means analysis underestimated the number of FC state transitions for G1, while the number of FC state transitions for G2 was recovered more accurately. Unless the correct value of *k* was estimated, Figure 7c shows that the combined analysis leads to artifactual differences in dFC between groups. One might think that fitting the two groups separately would solve this problem. Figure 7b demonstrates a modest improvement in the error in number of FC state transitions for G1 (yellow line) when *k* was underestimated, but a steep increase in error for G2 when *k* was overestimated. This deterioration when *k* > 6 is to be expected because the *k*-means algorithm had to identify more FC states than are actually present in the G2. We see in Figure 7d that for the separate analysis, incorrect group differences were again found when *k* was overestimated or grossly misspecified for either group.

**Figure 6:**
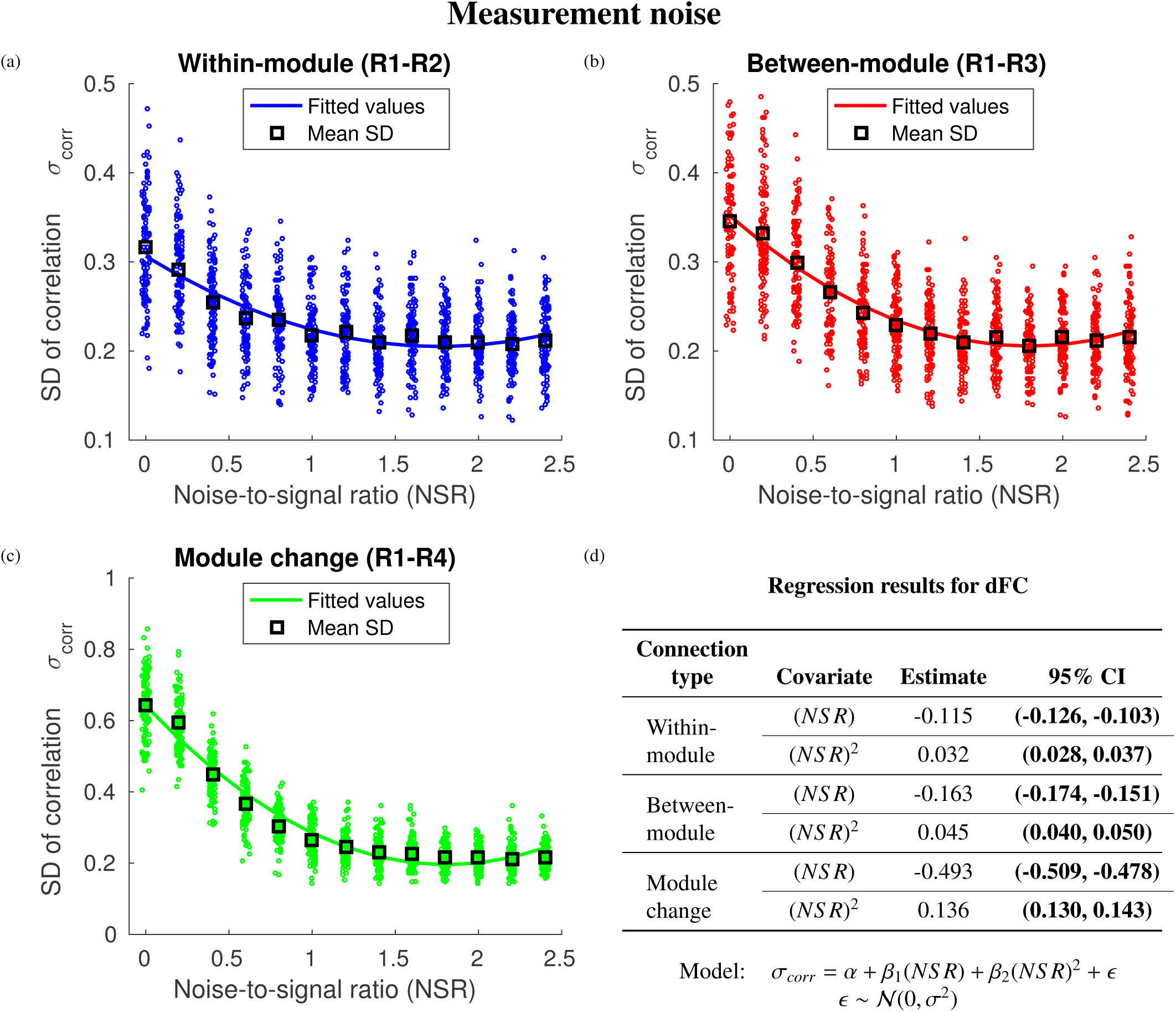
The impact of measurement noise on estimated dFC, measured by the SD of the correlation time series, between (a) statically connected regions (within-module), (b) unconnected regions (between-module), and (c) dynamically connected regions (module change), with (d) the results of a multiple regression. R_*i*_ refers to Region *i*. Region pair R1-R4 has a dynamic connection so the true dFC should be higher than region pairs R1-R2 and R1-R3, which have a static connection. Increased measurement noise resulted in decreased observed dFC for the three types of region pairs. This effect was particularly pronounced for the dynamically connected regions (module change). The multiple regression assesses the impact of measurement noise on estimated dFC with a statistically significant effect indicated by a 95% confidence interval (CI) in bold type. The solid lines in (a-c) correspond to the fitted values of the multiple regression.

Figures 7e and 7f illustrate the differences between the groups for both analyses in mean centroid error, which is a measure of how well the complete space-time connectivity structure is recovered. In this case, the combined analysis performed better for the young group than the old group when *k* was underestimated (*k* < 9), indicating that the recovered centroids were biased towards the 6 joint states. In contrast, if *k* was correctly specified or only slightly misspecified for either group, the separate analyses had a similar error in recovered connectivity structure. While these results suggest that the separate analyses did yield some improvement on the combined analysis, we caution that the problem of estimating *k* for both groups still needs to be addressed. As shown in Figure 7c and 7d, without an accurate estimation of *k*, one is likely to incorrectly infer the size of differences in dFC across groups, even for cases where there is no true difference. A common method for estimating the number of clusters, *k*, is the elbow plot shown in Figure 8. This demonstrates that it is not always straightforward to estimate *k* accurately: while we see a clear elbow for the separate analysis of G2 (purple dotted line), there is no obvious elbow for the separate analysis of G1 (yellow dotted line) or the combined analysis (solid black line).

**Figure 7:**
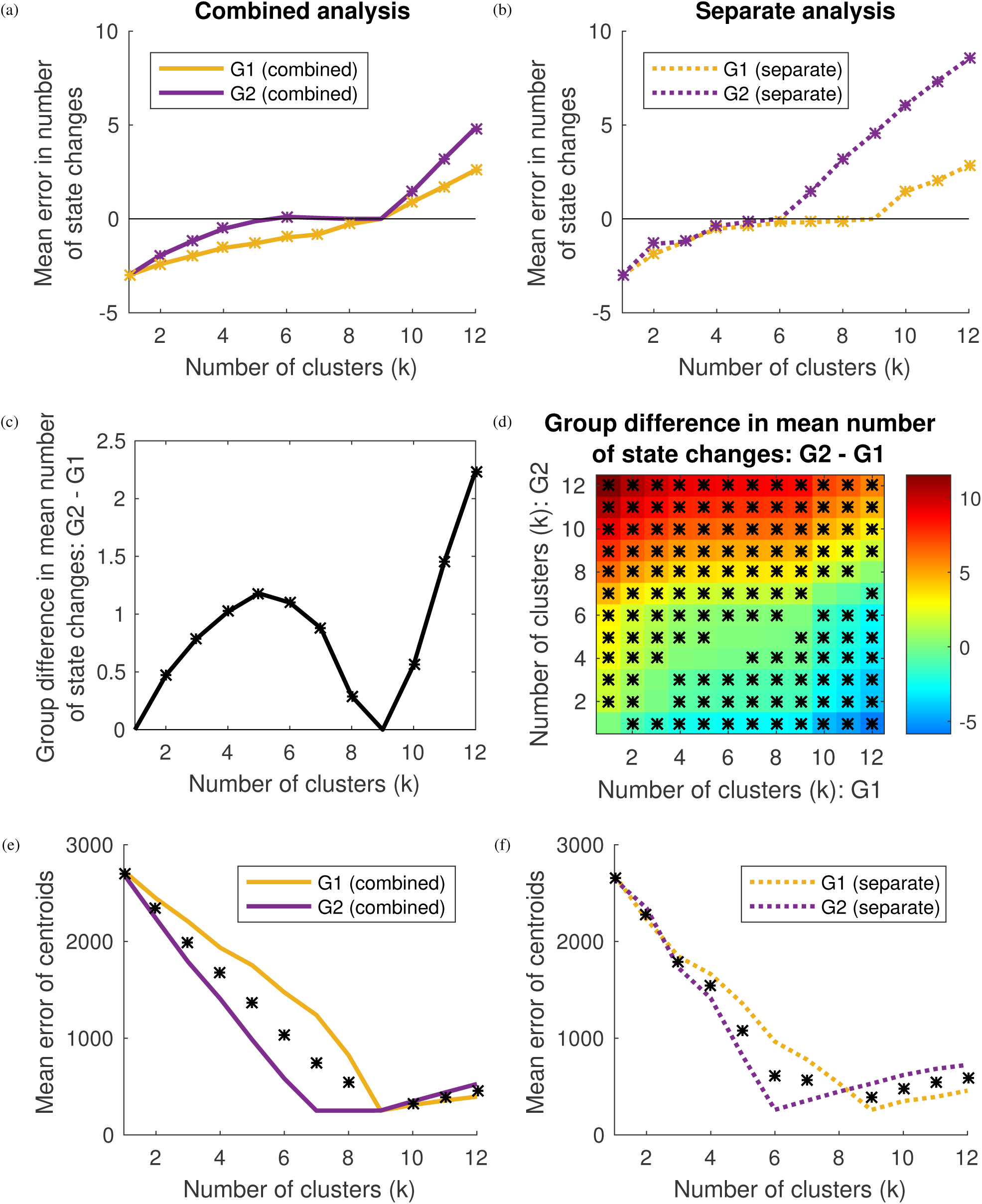
The true number of FC state transitions in this simulation is 3. Individuals in G1 could reach 9 FC states while individuals in G2 could reach only 6 of these 9 FC states. (a) When *k* < 9, the combined analyses underestimated the number of FC state transitions for G1 (solid yellow line), while the number of FC state transitions for G2 (solid purple line) was recovered more accurately. (b) The separate analysis showed an improvement in the error in number of FC state transitions for G1 (dotted yellow line) when *k* was underestimated. (c) Unless *k* was correctly estimated, the combined analysis yielded an incorrect group difference in number of FC state transitions. (d) For the separate analysis, incorrect group differences were only found when *k* was overestimated or grossly misspecified for either group. (e) In terms of recovered space-time connectivity structure, the combined analysis performed better for G2 (solid purple line) than G1 (solid yellow line) when k was underestimated. (f) If k was correctly specified or only slightly misspecified for either group, the separate analyses had a similar error in recovered connectivity structure. Asterisks in (a-d) indicate a statistically significant (*p* < 0.05) difference from zero, according to a Wilcoxon signed-rank test, while asterisks in (e-f) indicate a statistically significant (*p* < 0.05) difference between the two groups, according to a two-sample Wilcoxon rank-sum test.

**Figure 8:**
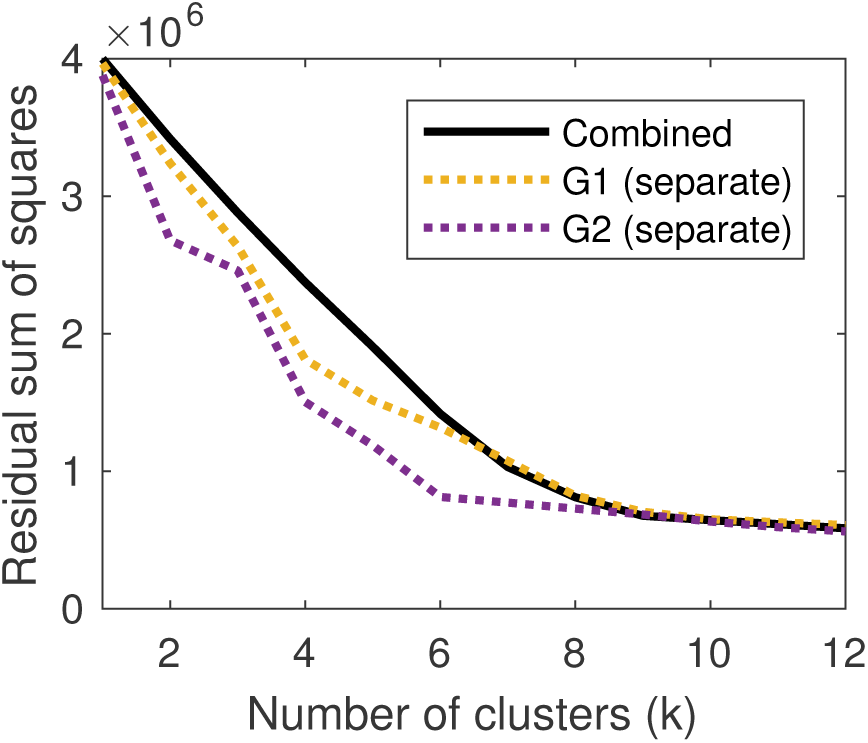
The RSS plots for the separate analysis of G2 has a clear elbow at *k* = 6 but there is no obvious elbow for the separate analysis of G1 or the combined analysis of both groups.

### 4.6. Multilayer modularity

While the *k*-means method represents an approach which aggregates data across subjects in order to glean information about functional connectivity dynamics, other methods analyse fMRI data on a subject-by-subject basis. For example, the multilayer modularity approach (Bassett et al. (2011)) characterises the correlation matrices for a single subject obtained from a sliding-window analysis as a multi-layered network. Each region in each window is assigned a module label by maximising a modularity index which depends on two hyperparameters *γ* and *ω* (see section 3.6 for details).

In this simulation, we again generated data for two groups of 50 individuals: those in G1 experienced one FC state transition while those in G2 experienced three FC state transitions. We applied the multilayer algorithm for all 100 individuals for *γ* = 0.75, 1, …, 2.5 and *ω* = 0.25, 0.5, …, 4. One measure of dFC in this approach is the mean “flexibility” of each brain region (see Methods). Thus, we now simulated a true group difference in the number of FC state transitions, and examined how accurately the FC states were recovered as a function of the methods hyperparameters. We also examined the error in the true vs estimated mean flexibility for each group.

Figure 9 illustrates the importance of parameter selection for the multilayer modularity approach. Figures 9a and 9c show that, in terms of the complete space-time connectivity dynamics, the optimal value for ω differed between the two groups. A lower value for ω generally resulted in more changes in module assignment across consecutive time windows. Since an individual in G1 experienced fewer FC state transitions, brain regions had fewer changes in module assignment across the course of the scan. Thus for G1, a higher value for ω was more effective in recovering the spatio-temporal connectivity structure.

We note that the optimal value for *γ* appears to be broadly the same for both groups. The parameter γ influences the resolution of the recovered network. A higher value for *γ* partitions the brain regions into more modules. Since ROIs were always partitioned into 5 modules for both groups, we would not expect the optimal value of *γ* to differ between the groups.

Figures 9b and 9d show that larger values of ω yield higher recovered mean flexibility for both G1 and G2. This effect, however, does not occur at the same rate for both groups. Figure 10 shows that different values of *γ* and *ω* result in different group differences in mean flexibility, to the extent that G1 is incorrectly found to be more flexible for some values of the hyperparameters. Note that the true difference in flexibility was approximately 0.14 (see Section 3.6) and this was not captured for any values of *γ* and *ω*. This suggests that caution should be taken when computing group differences, especially when an assumption of homogeneity is made.

## 5. Discussion

In this article, we have illustrated some of the limitations of current dFC methods when dealing with heterogeneity. We used a generic simulation framework to isolate various sources of heterogeneity, and showed that observed connectivity dynamics may be due to factors other than true changes in connectivity.

To investigate the effects of individual differences in neural autocorrelation, HRF shape, connectivity strength and measurement noise, we used the SD of correlation values across sliding windows as our measure of dFC. We calculated this measure for three types of connectivity: static, positively connected; static, unconnected; and dynamically connected (positively connected to unconnected). Increased neural autocorrelation resulted in higher dFC for statically connected regions but lower dFC for dynamically connected regions. A more temporally dispersed HRF produced higher dFC for all three connectivity types. In contrast, increased measurement noise yielded lower dFC across the three types of connectivity. Increased connectivity strength resulted in higher dFC for the dynamically connected regions but lower dFC for the positively statically connected regions. Together, these findings demonstrate that individual differences in dFC can be caused by various properties of the fMRI signal that are unrelated to the underlying neural connectivity dynamics.

We also demonstrated that common dFC methods may detect artifactual group differences in dynamic connectivity due to the assumptions that are made. For example, in a *k*-means analysis, it is often assumed that all individuals may attain the same set of FC states. If the hyperparameter *k* is incorrectly estimated, an incorrect group difference in the number of FC state transitions experienced may be detected if one group can attain more FC states than the other group. We note that these issues could in principle affect any FC state-based method which assumes homogeneity in attainable FC states across individuals.

**Figure 9:**
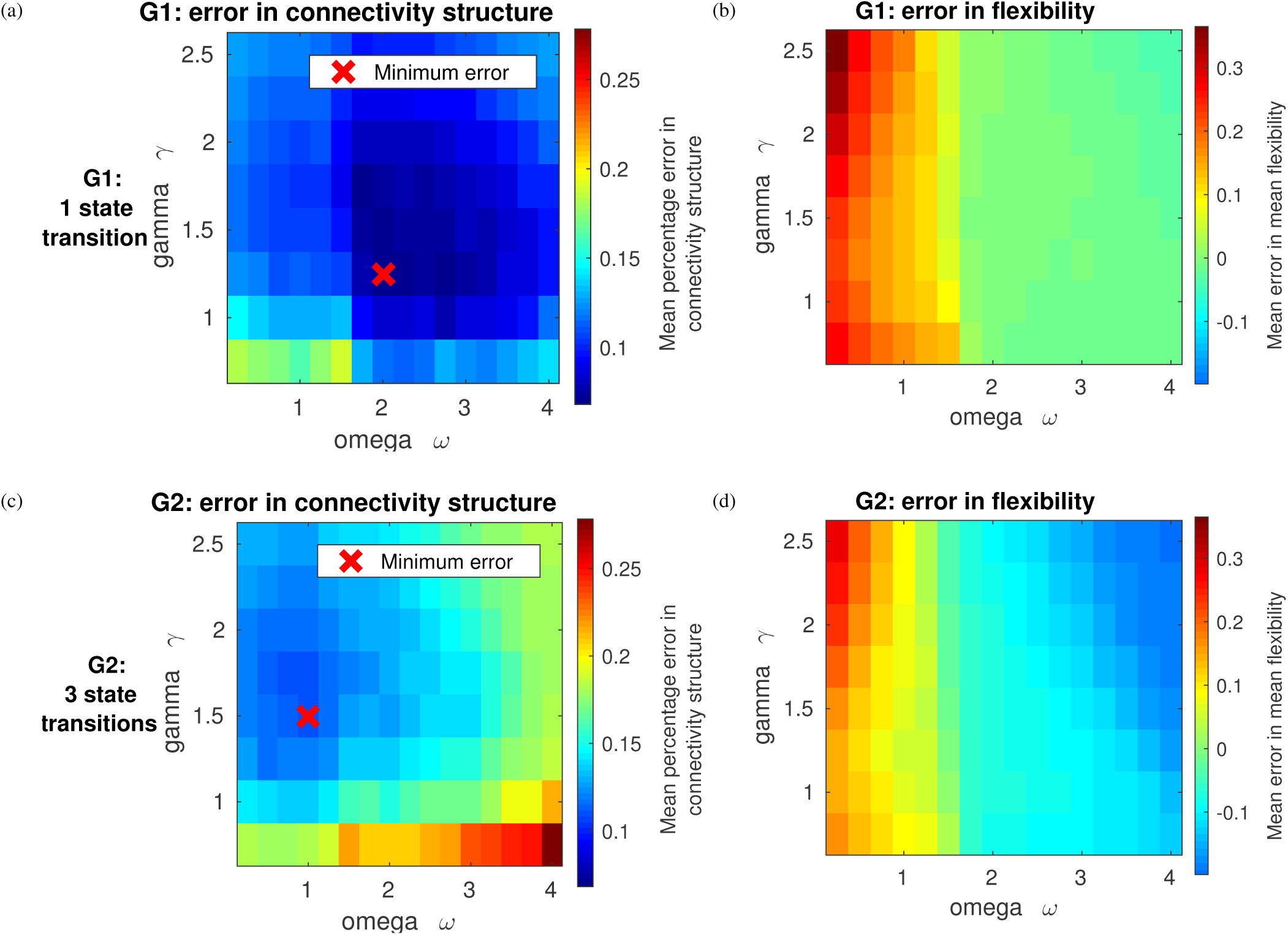
The effect of varying parameters *γ* and *ω* on performance of the multilayer modularity approach. Individuals in G1 experience 1 state transitions while individuals in G2 experienced 3 state transitions. Percentage error in connectivity structure is defined as the percentage of entries in the recovered incidence matrix that are equal to the corresponding entries of the true incidence matrix. A red cross indicates the pair of parameter values (*γ, ω*) which minimised the mean (across individuals) percentage error in connectivity structure. The flexibility of a region is defined as the number of times the region changes module assignment divided by the total possible number of module changes. (a) For G1, the optimal parameter values for recovering the connectivity structure were *γ* = 1.25, *ω* = 2. (b) Increasing *ω* resulted in decreased recovered flexibility for G1. (c) For G2, the optimal parameter values for recovering the connectivity structure were *γ* = 1.5, *ω* = 1. Thus, the optimal value for *ω* was markedly lower for G2 than for G1 while the optimal value for *γ* was slightly higher for G2 compared to G1. (d) Increasing *ω* also resulted in decreased recovered flexibility for G2, but at a greater rate than for G1.

More generally, care should be taken with any method that requires the selection of hyperparameters by the user. In particular, we demonstrated that group differences in mean flexibility detected by a multilayer modularity approach were regulated by the choice of hyperparameters. While one would expect individual-based methods such as the multilayer modularity approach to be more robust to heterogeneity, spurious group differences can nonetheless be found if hyperparameters are assumed constant across individuals.

One could attempt to optimise the choice of hyperparameters using the data. For example, in a k-means analysis, the number of clusters, k, could be estimated by a Variational Bayesian approach (Ghahramani et al. (1999)). For the multilayer modularity approach, one could use cross-validation across independent scans in an attempt to maximise stability of the recovered connectivity structure, though this assumes that dynamics are invariant across scans on the same individual. Alternatively, Bassett et al. (2013a) suggest choosing the values of *γ* and *ω* which yield connectivity structure that is most different from particular null models. Hyperparameter optimisation could be investigated in the future work, though our point is that such optimisation should allow for heterogeneity across individuals.

We focused on a number of likely sources of heterogeneity in fMRI signals, using effects of age to illustrate some of our examples. This is based on recent evidence of group differences in signal autocorrelation (Geerligs et al. (2016), Arbabshirani et al. (2014a)), HRF shape (Hanlon et al. (2016), Huettel et al. (2001), Aizenstein et al. (2004), D’Esposito et al. (1999)), and non-neural physiological noise levels (Geerligs et al. (2015), Mark et al. (2015)). Nevertheless, our findings apply in any situation where such heterogeneity may arise between individuals. Furthermore, there may be other sources of heterogeneity not investigated here that could have spurious effects on observed dFC. For example, we only considered variability in 2 out of the 7 HRF parameters; it is plausible that the remaining parameters also have an effect on estimated dFC. Similarly, we assumed that brain regions partition into 5 modules in each FC state, whereas it is conceivable that this could differ among individuals.

**Figure 10:**
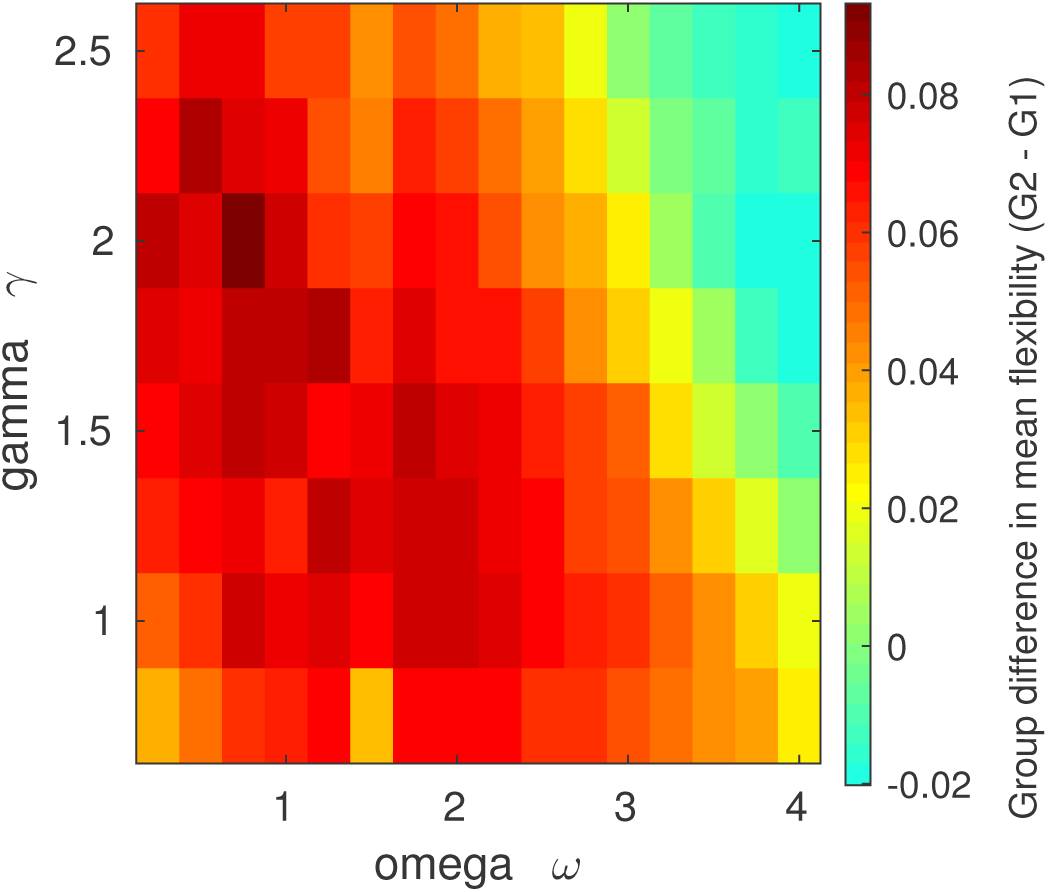
The effect of varying parameters *γ* and *ω* in the multilayer modularity approach on the recovered group difference in mean (across regions) flexibility. Individuals in G1 experience 1 state transitions while individuals in G2 experienced 3 state transitions. The flexibility of a region is defined as the number of times the region changes module assignment divided by the total possible number of module changes. Different values of *γ* and *ω* yield different group differences in mean flexibility. Note that the true difference here is 0.14, which is not attained for any values of the parameters.

We note that certain aspects of the simulation framework represent simplifications of the physical and physiological processes involved in fMRI neuroimaging. For example, the addition of white measurement noise is not realistic and noise related to head motion or vascular effects may have more regionally specific effects on connectivity estimates (Power et al. (2012)). Therefore the noise simulation should be interpreted as a cautionary result, and not as an illustration of effects of real noise sources in fMRI. We also assumed that the HRF is the same for all regions, although the shape of the HRF has been shown to vary from region to region (e.g. Schacter et al. (1997)). These assumptions, however, allowed us to isolate the impact of unaccounted heterogeneity. In particular, our current simulation framework had the distinct benefit of separating the underlying dFC structure from sources of heterogeneity such as neural autocorrelation, noise and HRF shape. We were thus able to manipulate dFC and these other sources of variation independently and show how observed dFC is affected. Further, in the Supplementary Material, we demonstrate that the effects of unrelated sources of variation persist across a range of parameter values, suggesting that these effects of dFC are not specific to this simulated dataset, but are in fact a more general phenomenon.

In this article, we do not address what drives these changes in functional connectivity. Recent observations suggest that dFC can be explained in terms of sparse brief events (Allan et al. (2015), Karahanoğlu and Van De Ville (2015), Liu and Duyn (2013), Tagliazucchi et al. (2012)). Our simulation framework is based on work by Allen et al. (2012) which attempts to find periods of recurrent patterns of functional connectivity, or FC states, across time and individuals. It may be that these periods are longer than the underlying neural processes due to the temporal limitations of fMRI. Many FC state-based studies work on the basis of certain assumptions about FC states regarding, for example, their discreteness and typical duration. However, little is actually known about the nature of FC states and future work is required to better understand how these neural processes drive observed FC states.

To illustrate the effects of heterogeneity in the number of attainable states and the number of state changes, we simulated binary differences between two groups of individuals. This represents a simplification since, in reality, it is likely that differences between individuals fall on a continuous spectrum and so we caution against dichotomising between groups. It should also be emphasised that while here we have isolated the impacts of different sources of heterogeneity, in reality they may appear in combination. Future work could investigate how different types of heterogeneity interact, or even counteract, to produce differences in observed dFC.

Although we have chosen to illustrate the above points with only a few methods, the issues should in general extend to other approaches. For example, we used Fisher-transformed Pearson correlation as our basic measure of functional connectivity because this is currently the most commonly used metric. Alternative connectivity measures, such as coherence or multiplication of temporal derivatives (Shine et al. (2015)), may be less susceptible to certain types of unaccounted heterogeneity: for example, it has been shown that coherence is robust against variability in the shape of the HRF between regions (Ashby (2011)). Nonetheless, the issues of hyperparameter selection in dFC methods still need to be addressed regardless of the connectivity metric used. With this aim, we provide the Matlab code used for the present simulations here: http://www.mrc-bsu.cam.ac.uk/software/miscellaneous-software/. We encourage interested readers to explore alternative metrics and dFC analysis methods.

## 6. Concluding remarks

We use simulated fMRI data to demonstrate the effect of various sources of heterogeneity on observed dFC. Our results show that individual differences in dFC may be due to non-dynamic features of the data. The choice of hyperparameters in common methods is also important: these are often assumed constant across individuals, which can result in spurious group differences in dFC. We recommend that future studies consider implicit assumptions of homogeneity in their analysis.

## Acknowledgements

The authors thank Steven M. Hill for his helpful comments on a draft of this manuscript. BL, SRW and RNH are supported by the UK Medical Research Council [Programme numbers U105292687 MC-A060-5PR10]. The Cambridge Centre for Ageing and Neuroscience (Cam-CAN) was support by the Biotechnology and Biological Sciences Research Council (grant number BB/H008217/1). LG is funded by a Rubicon grant from the Netherlands Organization for Scientific Research.

The Cam-CAN corporate author consists of the project principal personnel: Lorraine K Tyler, Carol Brayne, Edward T Bullmore, Andrew C Calder, Rhodri Cusack, Tim Dalgleish, John Duncan, Richard N Henson, Fiona E Matthews, William D Marslen-Wilson, James B Rowe, Meredith A Shafto; Research Associates: Karen Campbell, Teresa Cheung, Simon Davis, Linda Geerligs, Rogier Kievit, Anna McCarrey, Abdur Mustafa, Darren Price, David Samu, Jason R Taylor, Matthias Treder, Kamen Tsvetanov, Janna van Belle, Nitin Williams; Research Assistants: Lauren Bates, Tina Emery, Sharon Erzinlioglu, Andrew Gadie, Sofia Gerbase, Stanimira Georgieva, Claire Hanley, Beth Parkin, David Troy; Affiliated Personnel: Tibor Auer, Marta Correia, Lu Gao, Emma Green, Rafael Henriques; Research Interviewers: Jodie Allen, Gillian Amery, Liana Amunts, Anne Barcroft, Amanda Castle, Cheryl Dias, Jonathan Dowrick, Melissa Fair, Hayley Fisher, Anna Goulding, Adarsh Grewal, Geoff Hale, Andrew Hilton, Frances Johnson, Patricia Johnston, Thea Kavanagh-Williamson, Magdalena Kwasniewska, Alison McMinn, Kim Norman, Jessica Penrose, Fiona Roby, Diane Rowland, John Sargeant, Maggie Squire, Beth Stevens, Aldabra Stoddart, Cheryl Stone, Tracy Thompson, Ozlem Yazlik; and administrative staff: Dan Barnes, Marie Dixon, Jaya Hillman, Joanne Mitchell, Laura Villis.

